# Oligodendroglia generate vascular mural cells and neurons in the adult mouse brain

**DOI:** 10.1101/2023.09.12.549127

**Authors:** Qingting Yu, Kairan Yang, Zhouling Fan, Maojiao Huang, Ting Xu, Yanzhuo Yang, Zuisu Yang, Xiaosong He, Falei Yuan

## Abstract

**BACKGROUND:** Oligodendroglia encompass oligodendrocyte precursor cells (OPCs) and oligodendrocytes (OLs). In the grey matter of the cortex, nearly all OPCs divide slowly, yet they don’t differentiate solely into mature OLs, leaving the exact role of these OPCs in the grey matter enigmatic.

**METHODS:** Oligodendroglia, including OPCs, were traced using the Sox10 Cre-ER^T2^ reporter mice. We compared the effect of tamoxifen dissolved in different solvents on the fate of Sox10 cells. We also compared the effect of tamoxifen dosage on the fate of Sox10 cells. The differentiation of labeled red fluorescent protein (RFP) cells was analyzed using immunofluorescence staining.

**RESULTS:** Two groups of RFP cells, type A Sox10 (Sox10-A) cells, and type B Sox10 (Sox10-B) cells, were identified in the cortex, striatum, hippocampus, thalamus, and hypothalamus. Sox10-A cells differentiate into platelet-derived growth factor receptor-β (PDGFRβ)+, CD13+ pericytes, and smooth muscle myosin heavy chain 11 (MYH11) + smooth muscle cells when the mice received ethanol or high-dose tamoxifen. Sox10-B cells transform into glutamatergic neurons when the mice received high-dose tamoxifen. Sox10-B cells include perineurona OPCs and OLs.

**CONCLUSIONS:** This investigation provides evidence that a substantial proportion of oligodendroglia in the grey matter serve as mural cell precursors and neuronal precursors. These two phenomena may contribute to our understanding of neurodegenerative diseases.

**Graphic abstract:** 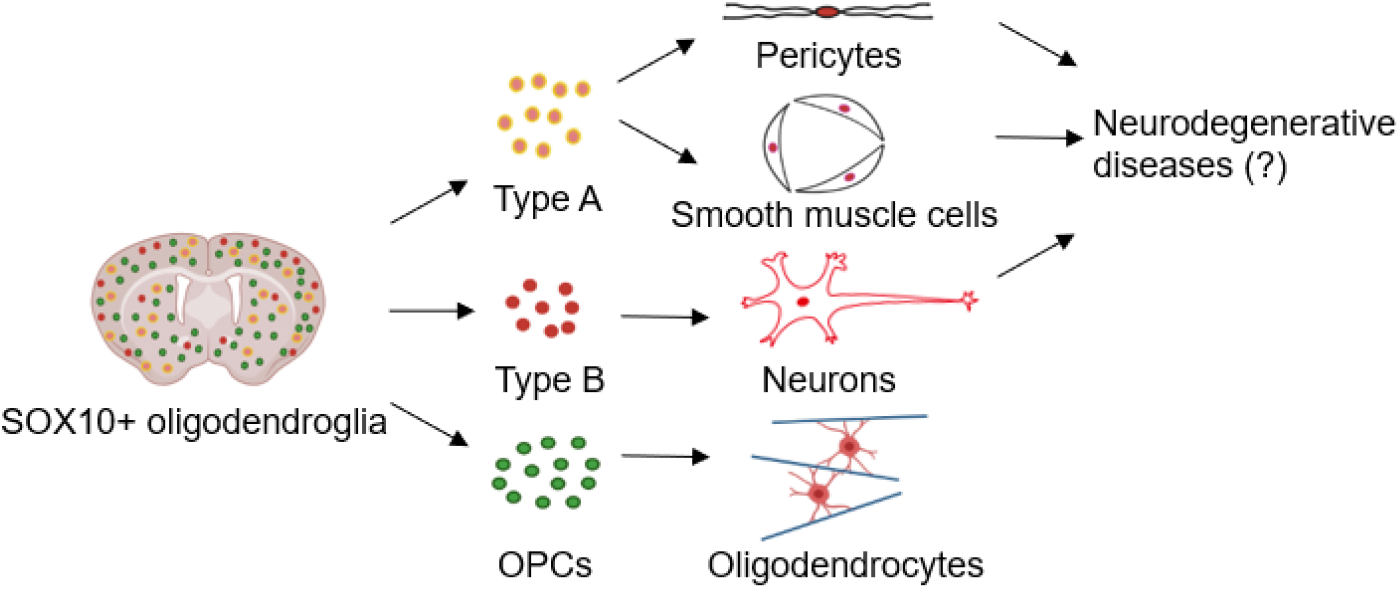

## 1. Introduction

Pericytes play a crucial role in maintaining the integrity of the blood-brain barrier (BBB) as a specialized group of perivascular cells in the brain. These cells closely associate with endothelial cells, covering the vasculature at a ratio ranging from 1:1 to 1:3 ^1^. Studies have demonstrated that pericytes are vital for the preservation of neuronal health, as their elimination has been shown to result in neuronal loss ^2^. Alzheimer’s disease (AD) has been speculated to have vascular origins, and researchers have observed significant alterations in capillaries near the pericyte soma in AD patients ^3^. Additionally, individuals with the apolipoprotein E4 (ApoE4) variant, known to increase the risk of AD, exhibit a lower abundance of pericytes ^4^. Gene expression studies have revealed that the mural cells of blood vessels, including pericytes, undergo the most substantial changes in AD patients ^5, 6^. In addition to their role in maintaining the BBB, pericytes are responsible for regulating brain blood flow and have been found to be more susceptible to cell death following ischemia compared to other neural cell types ^7^. Recent studies have also indicated that pericytes are the first responders during neuroinflammation, contrasting with the previously believed role of microglia ^8, 9^. Identifying pericytes is often accomplished by examining the expression of platelet-derived growth factor receptor β (PDGFRβ), cluster of differentiation 13 (CD13), and α-smooth muscle actin (α-SMA). However, the debate regarding whether brain pericytes express the contractile α-SMA still persists due to issues related to tissue fixation ^10^. It is worth noting that smooth muscle myosin heavy chain 11 (MYH11), previously considered specific to smooth muscle cells (SMCs), has also been discovered to be expressed in a specific subtype of pericytes ^11^. Currently, differentiating between SMCs and pericytes remains a challenge due to the lack of available methods ^12^.

Oligodendrocyte precursor cells (OPCs) represent the fourth type of glial cells in the brain. These slowly proliferating cells make up approximately 5% of all the neural cells in the brain. As the name implies, OPCs constantly differentiate into mature oligodendrocytes (OLs) ^13^. In recent years, OPCs have been found to be very active, like microglia ^14^. Neuron glia antigen-2 (NG2), platelet-derived growth factor receptor α (PDGFRα), oligodendrocyte lineage transcription factor 2 (Olig2) have been used for the identification of OPCs. They are also named as NG2 glia instead of NG2+ cells which include pericytes and vascular SMCs ^15, 16^. In white matter, OPC processes align parallel to axons, while in grey matter, they exhibit a radial morphology, similar to microglia ^17^. Interestingly, OPCs in the grey matter do not appear to differentiate solely into mature OLs ^18^. These cells are consistently found in close proximity to vascular endothelial cells ^19^, migrating along blood vessels ^20^, with some OPCs even establishing contact with nearby blood vessels through their processes ^21^. The debate surrounding whether OPCs differentiate into mature neurons has persisted for a considerable duration ^22-24^. In a recent paper, it was demonstrated that oligodendrocyte lineage cells can transfer nuclear and ribosomal material to neurons ^25^, a phenomenon believed to take place in healthy brains ^26^. Despite the progressive strides made in comprehending the diversity among OPCs, the exact functions and offspring of these cells continue to remain enigmatic.

Sex-determining region Y-related high mobility group-box 10 (Sox10) has been identified as being exclusively expressed in OPCs, myelinating OLs, and newly formed OLs in the adult mouse brain ^27^. It serves as a more specific marker for the oligodendrocyte lineage compared to NG2, PDGFRα, and Olig2 ^17^. Given the unknown physiological functions of oligodendrocyte lineage cells, an inducible Sox10-Cre tracing system was utilized in our study. This approach led us to discover a unique cluster of oligodendroglia in the mouse brain, which assumed the role of vascular mural cells under alcoholic conditions. Furthermore, our investigation demonstrated that the oligodendroglia-to-neuron conversion phenomenon observed in lineage tracing arises from tamoxifen toxicity, as opposed to being a healthy occurrence.

## 2. Methods and Materials

### 2.1 Animals

Sox10 Cre-ER^T2^ (#027651, Jackson Laboratory) and Ai9 (#007909, Jackson Laboratory) reporter mice were crossed, and male F1 generation at the age of 8 weeks were employed for induction of red fluorescent protein (RFP) using tamoxifen with various solvents, as specified in Results. Animal experiments were approved by the Experimental Animal Ethics Committee of Zhejiang Ocean University (#SCXK ZHE 2019-0031).

### 2.2 Immunofluorescence staining

All mice were sacrificed using carbon dioxide and subsequently underwent transcardial perfusion with saline, followed by 4% paraformaldehyde. The brain tissues were then fixed in 4% paraformaldehyde at 4°C for 1 h. Vibratome sectioning (ZQP-86, Zhisun Equipment Inc., Shanghai) was performed after embedding the brain tissues in low melting point agarose. Free-floating sections were permeabilized with 0.3% Triton X-100 and blocked with 1% bovine serum albumin in a 24-well cell culture plate. For immunofluorescence staining, the tissues were incubated overnight at 4°C with primary antibodies targeting specific proteins: CD13 (GTX75927, Genetex), Sox10 (AF2698, Beyotime), PDGFRβ (AF1042, R&D Systems; 14-1402-82, ThermoFisher), MYH11 (ab224804, Abcam), NG2 (ab5320, Millipore; ab129051, Abcam), aquaporin-4 (AQP4, 59678, CST), PDGFRα (558774, BD Biosciences), Ki67 (12202, CST; ab15580, Abcam), neuronal nuclei (NeuN, 266004, Synaptic Systems), microtubule-associated protein 2 (MAP2, 8707, CST), RFP (600-401-379, Rockland), Cre recombinase (MAB3120, Millipore), gamma-aminobutyric acid (GABA, A2052, Millipore), Reelin (ab312310, Abcam), the proto-oncogene c-Fos (2250, CST), doublecortin (DCX, 4604, CST), vesicular glutamate transporter 2 (VGLUT2, MAB5504, Millipore), adenomatous polyposis coli (APC, clone CC1, GTX16794, Genetex) and activating transcription factor 3 (ATF-3, HPA001562, Millipore). After three washes with PBS, immunofluorescence secondary antibodies (705-095-147, 112-095-003, 111-095-003, 115-095-003, 706-095-148 Jackson ImmunoResearch) were applied at room temperature for 1 h. Finally, the nucleus was counterstained with 4’,6-diamidino-2-phenylindole (DAPI) and the brain sections were imaged using a fluorescence microscope (Olympus BX41).

### 2.3 Statistical analyses

Statistical analysis was performed using GraphPad Prism 9.0. In all figures, data were represented as mean ± standard error of mean (SEM). Significance analysis using two-way analyses of variance (ANOVA) followed by Bonferroni post hoc test or unpaired t test. *P* < 0.05 was considered as statistical significance.

## 3. Results

### 3.1 Sox10 cells differentiate into pericytes and SMCs in the brains of adult mice

To investigate the role of oligodendrocyte lineage cells in the mouse brain, we employed Sox10 Cre-ER^T2^ reporter mice for fate mapping these cells. After two days of tamoxifen administration (intraperitoneal injection of tamoxifen dissolved in sunflower seed oil at a daily dose of 40 mg/kg), followed by a five-day interval, RFP+ oligodendroglia were found evenly distributed in the whole brain (Figure 1A). This group was designated as the 7D group. In a few brain sections, we observed the emergence of blood vessel-like RFP cells (1.85% ± 0.21%) in the cortex, which resembles vascular mural cells (n=4 mice, Figures 1B, C, L). These cells were identified as either pericytes (11.82% ± 0.78% PDGFRβ+ and 11.71% ± 1.06% CD13+) or SMCs (25.54% ± 2.03% MYH11+) (Figures 1D-F, M-O). Additionally, AQP4 staining was employed to identify BBB-covered capillaries, revealing that 82.73% ± 1.88% of the converted cells were pericytes of BBB (Figure 1G).

**Figure 1.**
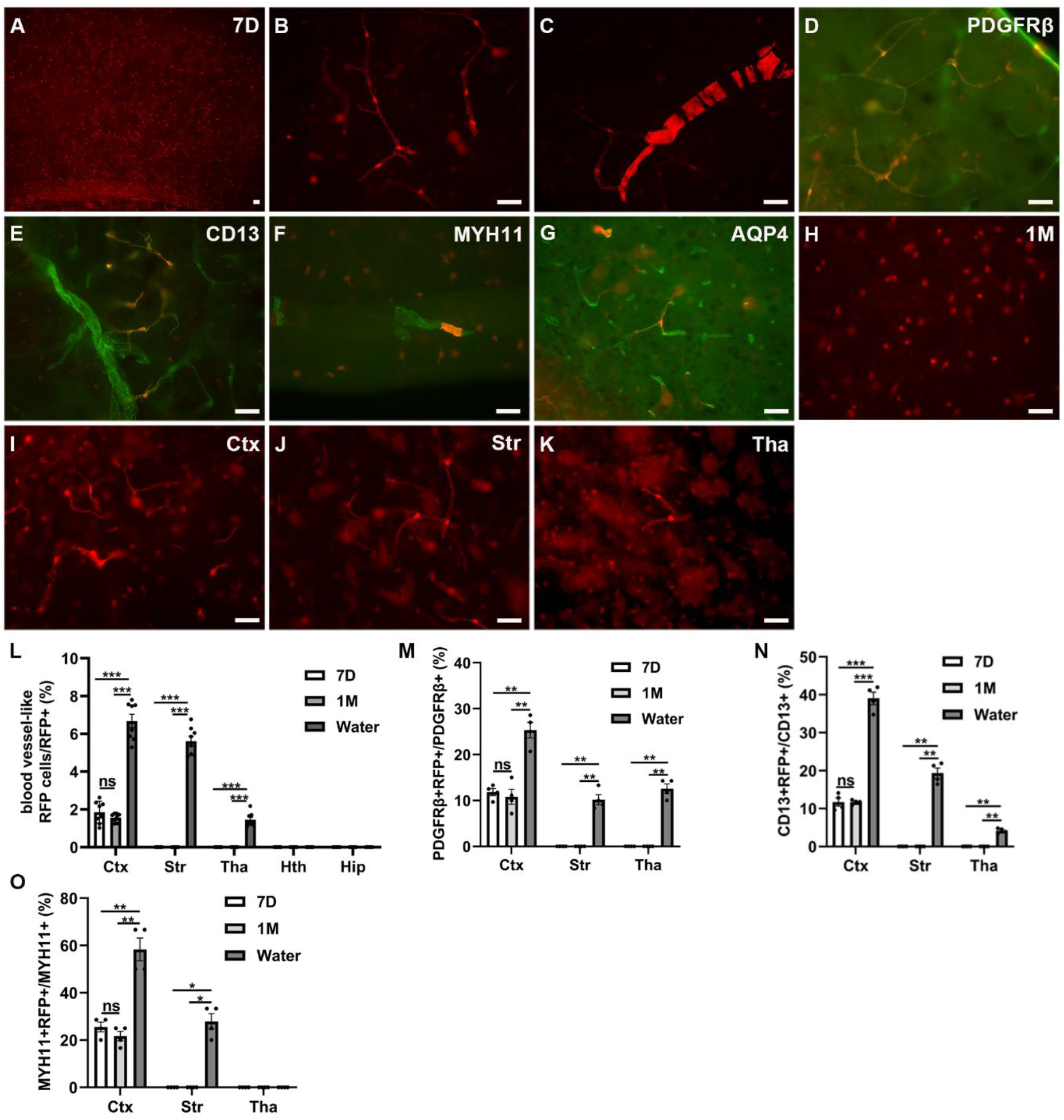
Fate-mapping of Sox10+ cells in the mouse brain. A small group of blood vessel-like RFP cells appeared in the 7D group (A-C), which was further confirmed by the immunofluorescent staining for PDGFRβ (D), CD13 (E), MYH11 (F), and AQP4 (G). RFP+ cells in the cortex of the 1M group (H). Blood vessel-like RFP cells in the cortex, striatum, and thalamus of the water group (I-K). Percentages of blood vessel-like RFP cells (L), PDGFRβ+ RFP cells (M), CD13+ RFP cells (N), MYH11+ RFP cells (O) in the different groups. Significance analysis using two-way analyses of variance (ANOVA) followed by Bonferroni post hoc test. Bars: 80μm for A, 20μm for B-K. The data are represented as mean ± SEM. \**p* < 0.05, \*\**p* < 0.01, \*\*\**p* < 0.001. Ctx, cortex; Str, striatum; Tha, thalamus; Hth, hypothalamus; Hip, hippocampus.

To confirm whether the small group of cells is continuously differentiating, mice were administered tamoxifen using the same strategy as in the 7D group, and the brain tissue was harvested 28 days later. This group was designated as the 1M group. We found that the 1M group had a percentage of blood vessel-like RFP cells of 1.56% ± 0.08%, along with 10.81% ± 1.63% PDGFRβ+, 11.69% ± 0.31% CD13+, and 21.67% ± 2.04% MYH11+ RFP cells, which were comparable to those in the 7D group (n=4 mice, Figures 1H, L-O). This indicated that the small amount of oligodendroglia-to-mural-cell conversion is not time course dependent.

We subsequently administered tamoxifen dissolved in ethanol via drinking water (50 mg of tamoxifen in 250mL of a 1% ethanol solution ^28^) and collected brain tissue 14 days later, referred to as the water group. Strikingly, the presence of RFP-converted mural cells increased significantly (6.68% ± 0.36% blood vessel-like RFP cells, 25.32% ± 1.71% PDGFRβ+, 39.04% ± 1.63% CD13+, and 58.33% ± 4.81% MYH11+) in the cortex of the water group (n=4 mice, Figures 1I, L-O; S1A-C). Furthermore, as shown in the Figures 1J, K, in the water group, these converted cells were also observed in the striatum (5.62% ± 0.26% blood vessel-like RFP cells, 10.19% ± 1.10% PDGFRβ+, 19.34% ± 1.36% CD13+, and 27.92% ± 3.29% MYH11+ RFP cells, Figures 1J, L-O) and the thalamus (1.45% ± 0.13% blood vessel-like RFP cells, 12.58% ± 1.05% PDGFRβ+, 4.18% ± 0.31% CD13+, Figures 1K-O). We conducted RFP staining and observed that these converted RFP cells were not artifacts in either the 7D group or the water group (Figures S1D, S1E). These Sox10+ RFP cells, possessing the capacity for oligodendroglia-to-mural-cell conversion, were designated as type A Sox10 (Sox10-A) cells.

### 3.2 A group of Sox10-A cells continue to express NG2, but not Sox10 or PDGFRα

As Sox10-A cells have the potential to differentiate into mural cells, we investigated whether these converted mural cells retain the expression of NG2, given that NG2 is frequently employed for OPC labeling and is sometimes used for labeling mural cells as well ^29^. We found that a part of converted Sox10-A cells maintained NG2 expression (32.42% ± 1.53% NG2+ RFP cells in the 7D group and 49.87% ± 2.16% in the water group, Figures 2A, B, K), while co-localization of Sox10 or PDGFRα expression with Sox10-A mural cells was not detected (Figures 2C-F). 4.28% ± 0.14% Ki67+ proliferating RFP cells were found in the cortex of the water group, and 1.70% ± 0.11% Ki67+ proliferating RFP cells were found in the cortex of the 7D group (arrows, Figures 2G, H, L). These results partially elucidate the reason why the glycoprotein NG2 is present on both OPCs and mural cells. This occurrence can be attributed to the fact that at least some Sox10-A cells continue to express NG2 even after differentiating into mural cells. Notably, mural cells expressed RFP but not Cre, providing evidence against the possibility of cell fusion (Figures 2I, 2J, S1D, S1E).

**Figure 2.**
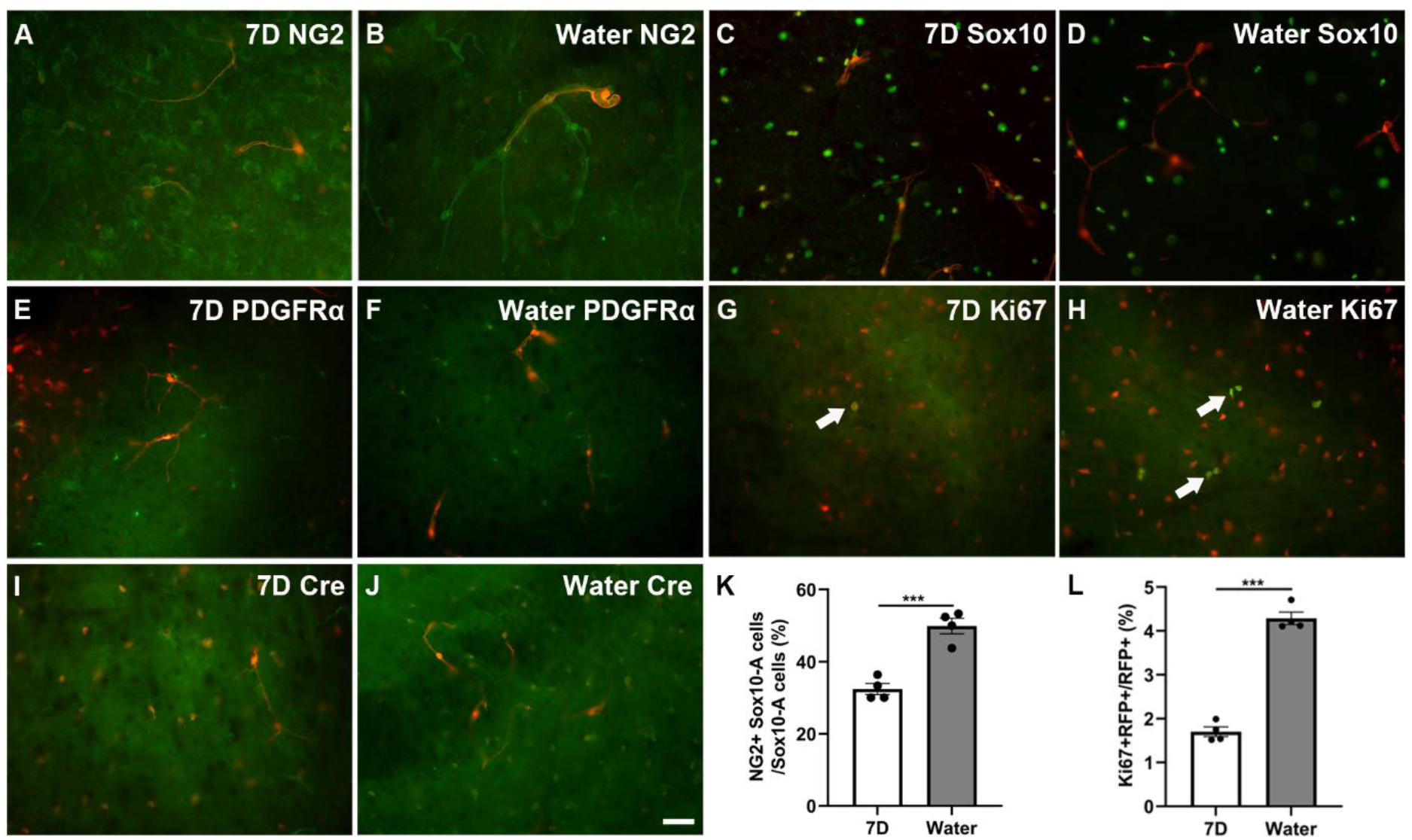
Representative images of the immunofluorescent staining for OPC related markers. Sox10-A cells co-localized with NG2 in the 7D group (A) and the water group (B), but did not co-localize with Sox10 (C, D) or PDGFRα (E, F). Ki67 and Cre immunostaining in the 7D group (G, I) and the water group (H, J). Percentages of NG2+ Sox10-A cells and proliferating cells in the different groups (K, L). The arrows depict Ki67+ RFP cells. Significance analysis using unpaired t test. Bar = 20μm. The data are represented as mean ± SEM. \*\*\**p* < 0.001.

### 3.3 Oligodendroglia-to-mural-cell conversion: influence of ethanol and tamoxifen

To determine whether oligodendroglia-to-mural-cell conversion is associated with ethanol exposure, we administered tamoxifen to mice using either pure sunflower oil or a mixture of sunflower oil and ethanol in a ratio of 9:1 via gavage (40 mg/kg per day) for three days. Subsequently, the mice were sacrificed 7 days after the initial tamoxifen dosage. The two respective groups were designated as the oil group and the oil/ethanol group.

In the oil group, Sox10-A cells constituted 1.46% ± 0.13% of all RFP cells (accounting for 10.62% ± 0.58% CD13+ pericytes and 23.06% ± 1.21% MYH11+ SMCs). This proportion was significantly higher in the oil/ethanol group, where Sox10-A cells represented 2.90% ± 0.16% of all RFP cells (accounting for 18.89% ± 0.49% CD13+ pericytes and 33.54% ± 1.54% MYH11+ SMCs), and this distribution was exclusively observed in the cortex (n=4 mice, Figure 3A, B, D-F, S2A, S2B, S2D, S2E). These findings indicate that ethanol facilitates oligodendroglia-to-mural-cell conversion.

**Figure 3.**
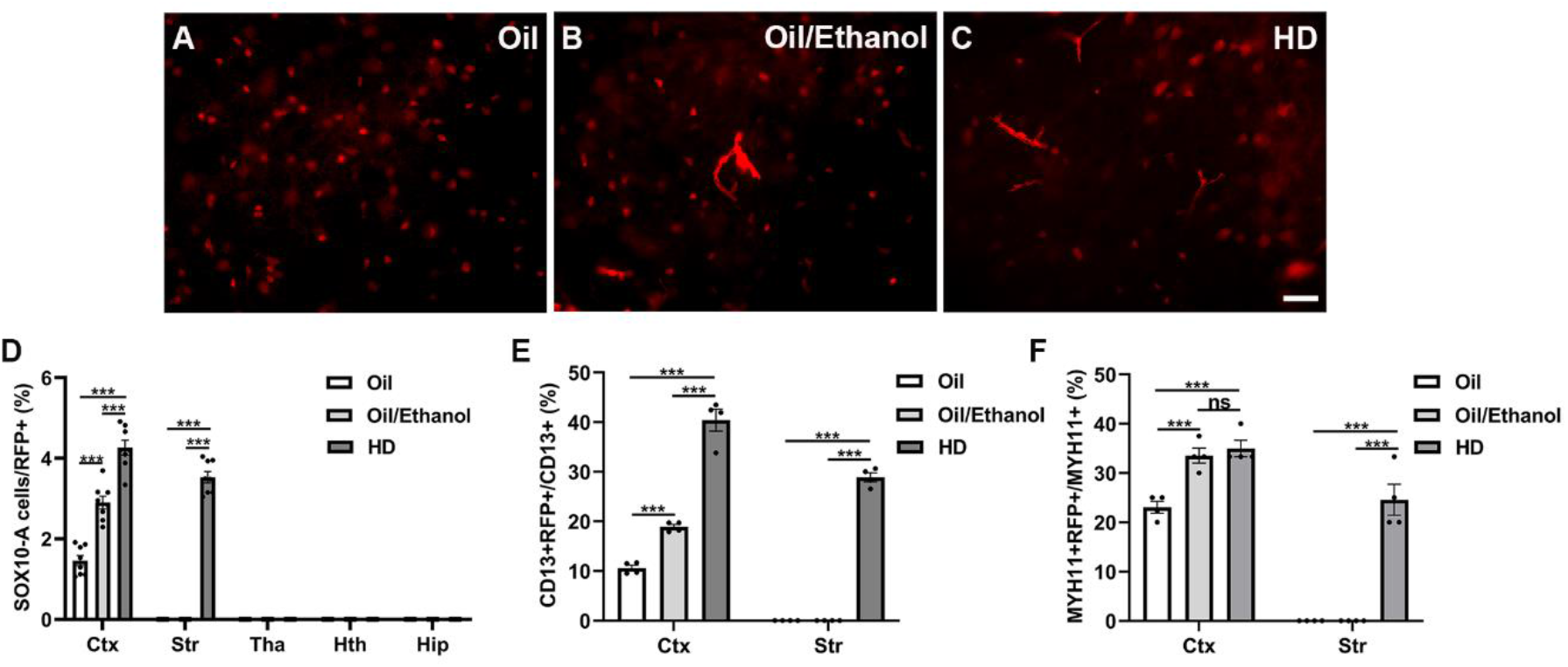
Oligodendroglia-to-mural-cell conversion in the three groups. Representative images of Sox10-A cells in the cortex of the oil group (A), the oil/ethanol group (B), and the high-dose group (C). Percentages of Sox10-A cells (D), CD13+ Sox10-A cells (E), MYH11+ Sox10-A cells (F) in the three groups. Significance analysis using two-way analyses of variance (ANOVA) followed by Bonferroni post hoc test. Bar = 20μm. The data are represented as mean ± SEM. \*\*\**p* < 0.001. Ctx, cortex; Str, striatum; Tha, thalamus; Hth, hypothalamus; Hip, hippocampus; HD, high dose.

In addition, the mice received high doses of tamoxifen via gavage (160 mg/kg daily, sunflower seed oil: ethanol= 9: 1) for three days, and brain tissue was harvested four days later ^30^. This group was designated as the high-dose group. Oligodendroglia-to-mural-cell conversion was significantly increased compared with the oil group and the oil/ethanol group, suggesting that high doses of tamoxifen also contributes to the conversion (4.26% ± 0.18% Sox10-A cells, 40.42% ± 2.22% CD13+ pericytes, and 35.00% ± 1.67% MYH11+ SMCs) (n=4 mice, Figures 3C-F, S2C, S2F). And these converted cells were also observed in the striatum (3.53% ± 0.14% Sox10-A cells, 28.89% ± 0.91% CD13+ pericytes, and 24.58% ± 3.15% MYH11+ SMCs, Figures 3D-F).

### 3.4 Oligodendroglia-to-neuron conversion is associated with tamoxifen dosage

Unexpectedly, RFP cells displaying large cell bodies and long projections were observed in the layer 2/3 of the cortex (7.93% ± 0.37% of RFP cells) in the high-dose group, as well as in the piriform cortex where the OPC-to-neuron phenomenon has previously been documented (Figures 4A, S3A) ^22^. These cells were also found in the granule cells of the dentate gyrus (DG), striatum, thalamus, and hypothalamus of the high-dose group, accounting for 3.77% ± 0.20%, 0.25% ± 0.01%, 0.35% ± 0.05%, 0.28% ± 0.02% of RFP cells, respectively. Notably, these cells exhibited neuronal markers NeuN and MAP2 (n=4 mice, arrows, Figures 4A-C, O). Conversely, RFP+ neurons were absent in the oil group (referred to as the low-dose group for convenience) and also in the 1M group (Figures 4D, E, S3B). RFP expression was confirmed in the RFP cells of the low-dose group and in the RFP neurons of the high-dose group (Figures 4F, G). These Sox10+ RFP cells, possessing the capacity for oligodendroglia-to-neuron conversion, were designated as type B Sox10 (Sox10-B) cells.

**Figure 4.**
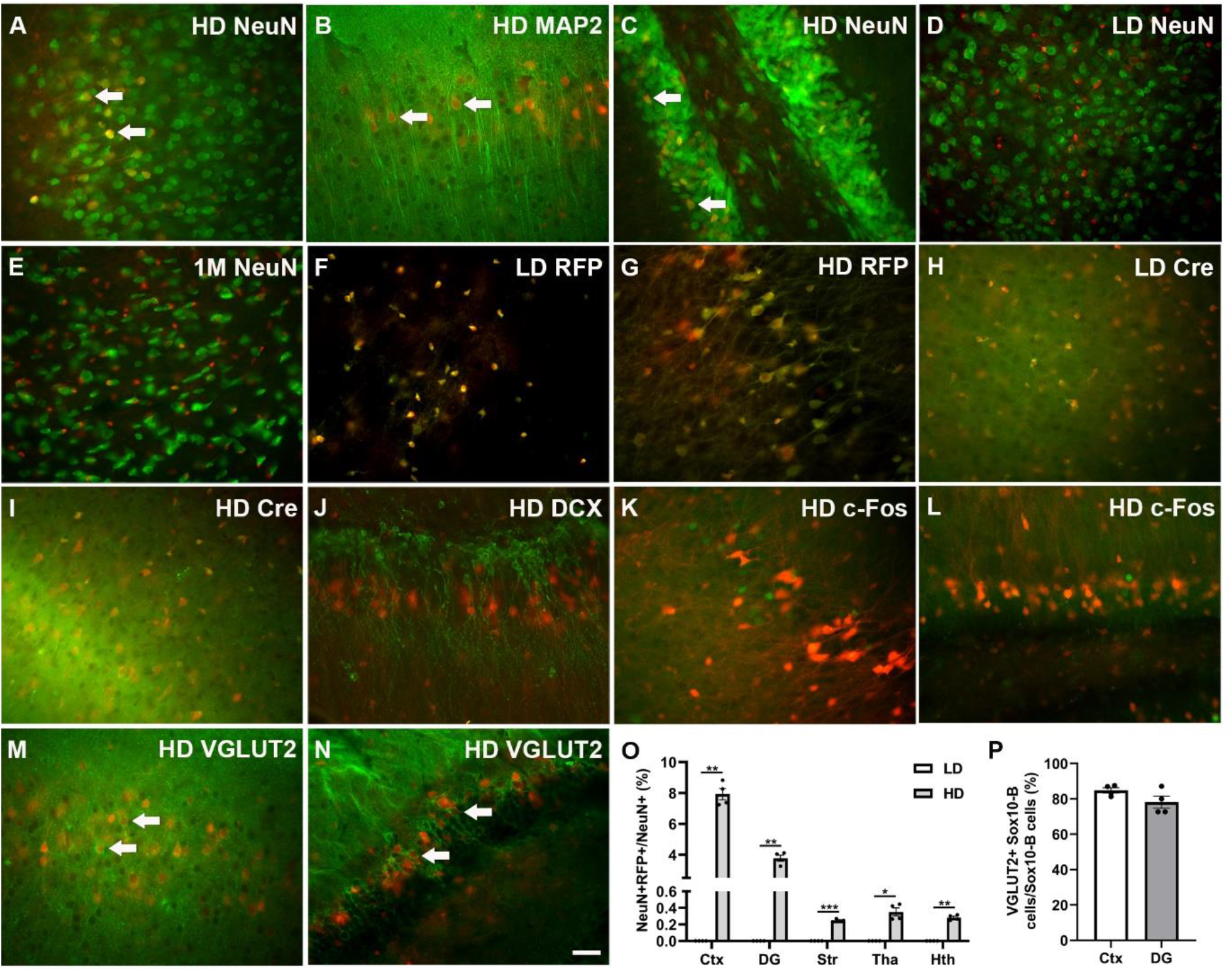
Sox10-B cells transform into neurons. Representative images of immunostaining for NeuN and MAP2 in layer 2/3 of the cortex of the high-dose group (A, B). NeuN immunostaining results in the DG of the high-dose group (C), in the cortex of the low-dose group (D), and in the cortex of the 1M group (E). Expressions of RFP (F, G) and Cre (H, I) in the low-dose group and high-dose group. Immunostaining for DCX (J), c-Fos (L), VGLUT2 (N) in the DG, and for c-Fos (K), VGLUT2 (M) in the cortex of the high-dose group. The percentage of RFP+ neurons in the low-dose and high-dose groups (O). The percentage of VGLUT2+ Sox10-B cells in the cortex and DG of the high-dose group (P). The arrows indicate Sox10-B cells. Significance analysis conducted using two-way analyses of variance (ANOVA) followed by Bonferroni post hoc test or unpaired t test. Bar = 20μm. The data are presented as mean ± SEM. \**p* < 0.05, \*\**p* < 0.01, \*\*\**p* < 0.001. Ctx, cortex; Str, striatum; Tha, thalamus; Hth, hypothalamus; LD, low dose; HD, high dose.

The Cre protein was identified in RFP cells, with no presence of RFP neurons immunoreactive for Cre. This observation indicates that the oligodendroglia-to-neuron phenomenon is not the result of cell fusion (Figures 4H, I, S3C-F). The underlying cause, whether direct differentiation or cell material transfer, requires further investigation. Additionally, to explore the possible connection of the phenomenon with neural stem cells, immunostaining for DCX was conducted. RFP neurons did not show co-localization with DCX in the DG or piriform cortex (Figures 4J, S3G), indicating that Sox10-B cells may not be associated with stem cell-related neurogenesis. The formation of Sox10-B neurons is not associated with neuronal excitation, as validated by c-Fos staining (Figures 4K, L). These neurons were predominantly identified as glutamatergic, positive for VGLUT2 in the cortex (84.76% ± 1.49%) and DG (78.14% ± 3.44%), but negative for GABA or Reelin (arrows, Figures 4M, N, P, S3H-K). Therefore, Sox10-B neurons are pyramidal neurons in the cortex or granule neurons in the DG.

### 3.5 Sox10-B cells: neuronal satellite oligodendroglia

Satellite oligodendroglia occupy specific positions and maintain close contact with neurons ^25, 31^. The proportion of satellite oligodendroglia was 24.14% ± 1.75% and 11.11% ± 1.24% in the cortex, 15.14% ± 1.15% and 5.26% ± 0.37% in the DG, 23.80% ± 1.25% and 16.69% ± 0.31% in the striatum, 15.96% ± 1.06% and 13.85% ± 0.65% in the thalamus, 20.14% ± 1.62% and 14.28% ± 1.11% in the hypothalamus of the low-dose group and the high-dose group, respectively (arrows, Figures 5A-D, I). The number of PDGFRα+ satellite RFP cells was 28 pairs in the low-dose group and 8 pairs and the high-dose group (Figures 5J). The number of CC1+ satellite RFP cells was 22 pairs in the low-dose group and 7 pairs and the high-dose group (Figures 5K). This indicates that Sox10-B cells include OPCs and OLs.

**Figure 5.**
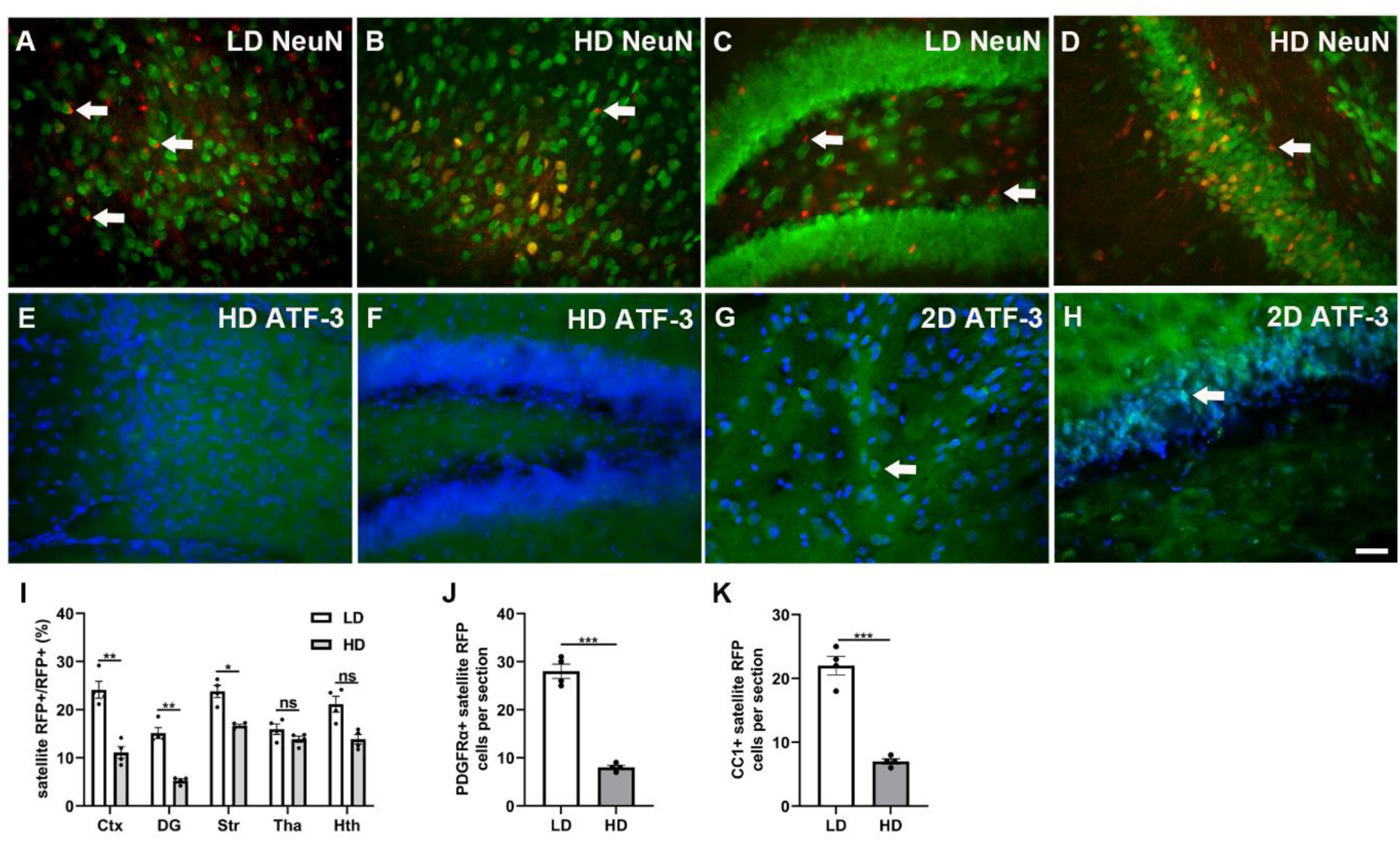
Neuronal satellite oligodendroglia in the mouse brain. Representative images of satellite oligodendroglia in layer 2/3 of the cortex and the DG in the low-dose group and high-dose group (arrows, A-D). Immunostaining for ATF-3 in layer 2/3 of the cortex and in the DG in the high-dose group and the 2D group (E-H). Arrows indicate ATF3+ neurons. Percentages of satellite oligodendroglia in the low-dose and high-dose group (I). Percentages of PDGFRα+ OPCs and CC-1+ OLs in the low-dose and high-dose group (J, K). Significance analysis using two-way analyses of variance (ANOVA) followed by Bonferroni post hoc test or unpaired t test. Bar = 20μm. The data are represented as mean ± SEM. \**p* < 0.05, \*\**p* < 0.01, \*\*\**p* < 0.001. Ctx, cortex; Str, striatum; Tha, thalamus; Hth, hypothalamus, LD, low dose; HD, high dose.

To assess whether oligodendroglia-to-neuron conversion involves neuronal stress, we conducted ATF-3 immunostaining, a protein related to cellular stress, and found that it was not expressed in the neurons in the cortex or DG of the high-dose group (Figure 5E, F). Subsequently, we administered a single dose of tamoxifen (160 mg/kg, sunflower oil: ethanol = 9:1), and mouse brains were harvested 48 hours later, designated as the 2D group. High levels of ATF-3 expression were observed in the pyramidal neurons of the cortex and granule cells of the DG (indicated by arrows, Figure 5G, H). Therefore, the initiation of neuronal injury and the absence of satellite oligodendroglia appear to be linked to oligodendroglia-to-neuron conversion, and these missing satellite oligodendroglia could be Sox10-B cells.

## 4. Discussion

In this study, we have identified two distinct populations of Sox10 cells: one comprising mural cell precursors called Sox10-A cells, which have the potential to transform into vascular mural cells, and another consisting of cells termed Sox10-B cells, which can differentiate into pyramidal neurons in the cortex and granule neurons in the DG. The transition of Sox10-A cells to mural cells is attributed to the use of ethanol as a solvent for tamoxifen and the dosage of tamoxifen. The transformation of Sox10-B cells into neurons is associated with the toxicity of tamoxifen, particularly at high dosages.

The substitution of pericytes in this study may be attributed to pericyte injury ^32^. In our pilot study, wild-type mice were intraperitoneally injected with alcohol for 30 minutes, and during this period, some pericytes in the cortex were labeled with intravenously injected propidium iodide (data not shown). This labeling indicated the cell membrane leakage of pericytes after alcohol treatment. In a previous study, we observed inflammation-related autofluorescence in nearly all major brain cells, including endothelial cells, but not in pericytes ^33^. Considering that autofluorescence tends to increase over time, this phenomenon could be attributed to the vulnerability of pericytes.

Furthermore, our findings highlight the unique presence of OPCs as a characteristic component of mural cells during brain angiogenesis ^34^, as well as the common occurrence of bipolar OPCs in diseased conditions, a factor that has received relatively less attention in the context of mural cells ^35^. Importantly, in the brains of AD patients, while astrocytes and microglia do not display signs of aging, OPCs exhibit aging characteristics ^36^. Additionally, a recent study has indicated that damage to oligodendroglia is an early event in the development of AD ^37^.

NG2 is commonly employed as a marker for identifying OPCs, but it is also occasionally used for identifying mural cells ^38^. In our observations, we noticed that not all converted Sox10-A cells were positive for NG2. This variability could be due to the challenges associated with NG2 immunofluorescent staining using different fixation methods ^39^. Relying solely on Sox10 fate mapping is inadequate for distinguishing between myelinating OLs and OPCs. The theory of new mural cell formation from myelinating OLs appears implausible, as myelinating OLs exhibit remarkable stability over time ^40^. The notion of OLs transforming into mural cells seems improbable as mature OLs are postmitotic cells that do not express NG2, while some converted Sox10-A cells still maintain NG2 expression ^41^.

Our observation aligns with the hypothesis that there exists a distinct population of OPCs in the grey matter that do not participate in oligodendrocyte development ^18^. To emphasize the morphological differences from conventional OPCs, some researchers have introduced terms such as polydendrocytes ^42^, or synantocytes ^43^, which resemble Sox10-A cells.

The replacement of neurons may be ascribed to neuronal injury, as indicated by the presence of the cell stress-related protein ATF-3 in neurons following tamoxifen administration ^44^. OPCs establish synapses with both glutamatergic and GABAergic neurons ^45^. One immediate question arises: why does the transformation of Sox10 cells into neurons exclusively occur in glutamatergic neurons rather than GABAergic neurons? Interestingly, immunostaining for c-Fos showed that the converted neurons may not be associated with neural excitation.

Neither of these phenomena seems to result from cell fusion, as Cre was not detected in the converted mural cells or neurons. The presence of RFP was further validated through immunofluorescence staining of RFP, which also eliminates the possibility of artifacts caused by autofluorescence. Additionally, cells may transfer both proteins and RNA through tunneling nanotubes ^46^. Further research is required to determine whether Sox10-A mural cells and Sox10-B neurons result from direct differentiation or merely a transfer of RFP from oligodendroglia to neurons.

Taken together, we have discovered two groups of Sox10 cells that can transform into vascular mural cells and neurons when subjected to injury. In our future studies, we will delve into the connection between their inhibition and neurodegeneration.

## Author contributions

Conceptualization: FLY, XSH; Data curation: QTY, FLY, KRY, MJH; Formal analysis: QTY, FLY; Funding acquisition: FLY; Investigation: QTY, XSH, MJH, FLY, KRY, ZLF, TX, YZY, ZSY; Methodology: QTY, FLY; Writing-original draft: QTY, KRY, FLY; Writing-review & editing: FLY.

