## Supplemental Figures for "Oligodendroglia generate vascular mural cells and neurons in the adult mouse brain"

**Supplemental data**

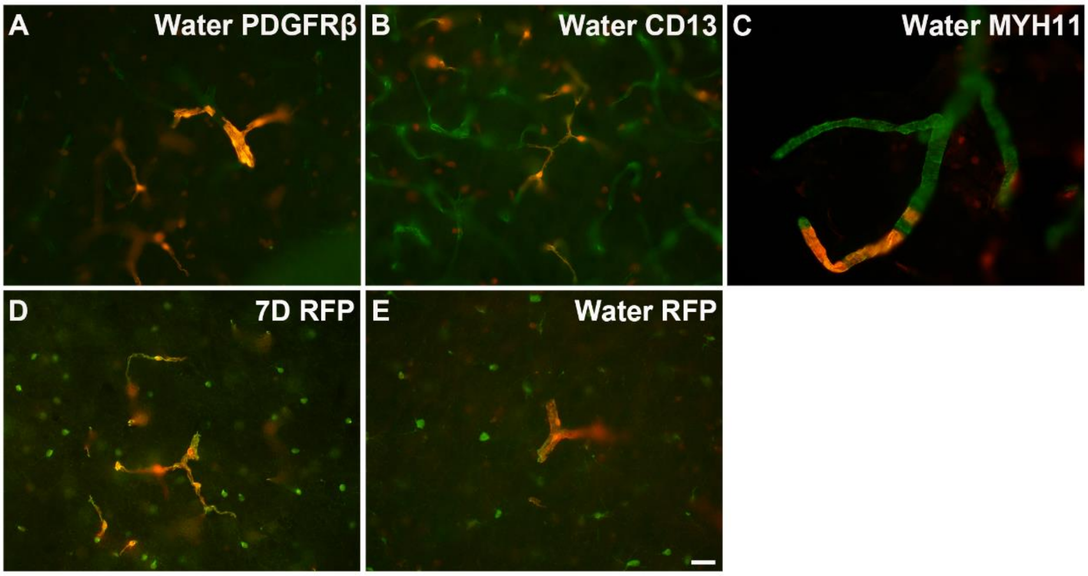


**Figure S1. Oligodendroglia-to-mural-cell conversion in the water group and 7D group.** Representative images of the immunostaining for PDGFRβ, CD13 and MYH11 in the cortex of the water group (A-C). RFP immunostaining in the cortex of the 7D group and water group (D, E). Bar = 20μm.

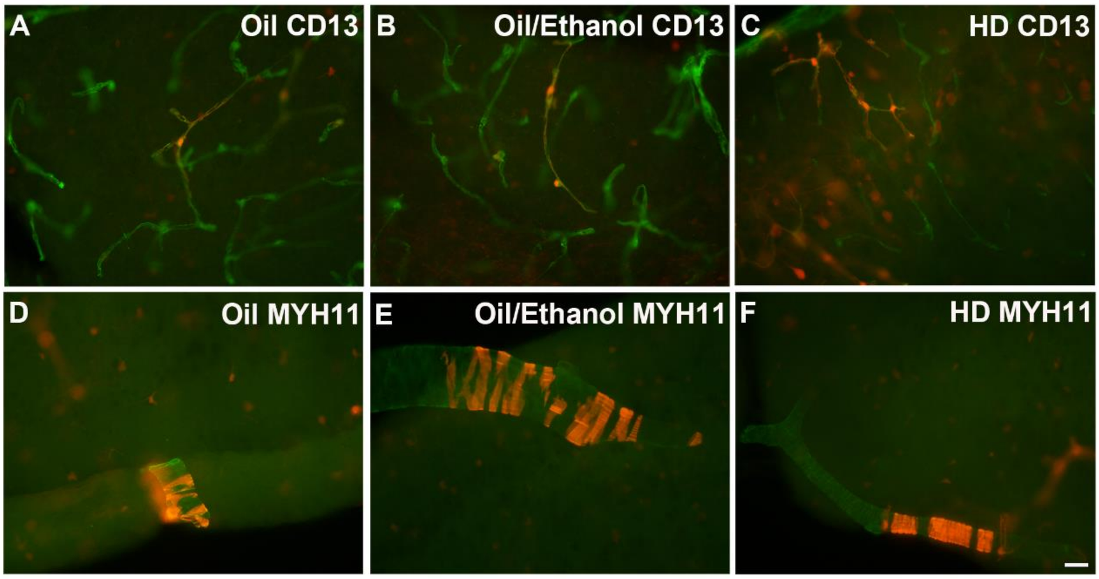


**Figure S2. Representative immunostaining images of mural cell proteins.**

Sox10-A cells were observed in the oil group, the oil/ethanol group, and the high-dose group, as confirmed by the immunofluorescent staining for CD13 (A-C) and MYH11 (D-F) in the cortex. Bar = 20μm. HD, high dose.

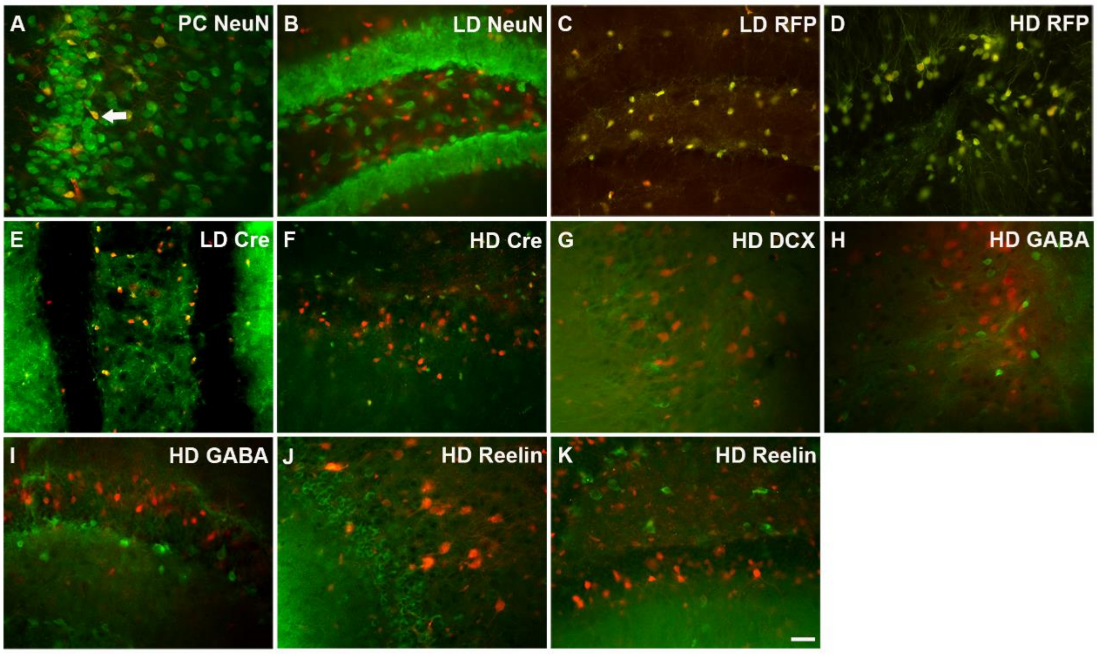


**Figure S3. Representative immunostaining images for identifying oligodendroglia-to-neuron conversion**. NeuN immunostaining in the piriform cortex of the high-dose group (A) and in the DG of the low-dose group (B). The arrow indicates a NeuN+ RFP cell. Expressions of RFP and Cre in the DG of the low-dose group and high-dose group (C-F). Immunostaining for DCX in the cortex of the high-dose group (G). Sox10-B neurons did not colocalize with GABA (H, I) or Reelin (J, K) in the cortex or DG of the high-dose group. Bar = 20μm. LD, low dose; HD, high dose; PC, piriform cortex; DG, dentate gyrus.
